# Investigations into hydrogen sulfide-induced suppression of neuronal activity in vivo and calcium dysregulation in vitro

**DOI:** 10.1101/2022.11.14.516514

**Authors:** Dong-Suk Kim, Isaac N. Pessah, Cristina M Santana, Benton Purnell, Rui Li, Gordon F Buchanan, Wilson K. Rumbeiha

**Author notes:** Wilson K. Rumbeiha, 1020 VM3B, 1089 Veterinary Medicine Dr. Davis, CA 95616.

## Abstract

Acute exposure to high concentrations of hydrogen sulfide (H_2_S) leads to sudden death and, if survived, lingering neurological disorders. Clinical signs include seizures, loss of consciousness, and dyspnea. The proximate mechanisms underlying H_2_S-induced acute toxicity and death have not been clearly elucidated. We investigated electrocerebral, cardiac and respiratory activity during H_2_S exposure using EEG, EKG and plethysmography. H_2_S suppressed electrocerebral activity and disrupted breathing. Cardiac activity was comparatively less affected. To test whether Ca^2+^ dysregulation contributes to H_2_S-induced EEG suppression, we developed an in vitro real-time rapid throughput assay measuring patterns of spontaneous synchronized Ca^2+^ oscillations in cultured primary cortical neuronal (PCN) networks loaded with the indicator Fluo-4 using the fluorescent imaging plate reader (FLIPR-Tetra^®^). Sulfide >5 ppm dysregulated SCO patterns in a dose-dependent manner. Inhibitors of NMDA and AMPA receptors magnified H_2_S-induced SCO suppression. Inhibitors of L-type voltage gated Ca^2+^ channels (VGCC) and transient receptor potential (TRP) channels prevented H_2_S-induced SCO suppression. Inhibitors of T-type VGCC, ryanodine receptors, and sodium channels had no measurable influence on H_2_S-induced SCO suppression. Exposures to >5 ppm sulfide also suppressed neuronal electrical activity in PCN measured by multi-electrode array (MEA), an effect alleviated by pretreatment with the nonselective TRP inhibitor 2-APB. The TRP inhibitor also reduced PCN cell death from sulfide exposure. These results improve our understanding of the role of different Ca^2+^ channels in acute H_2_S-induced neurotoxicity and identify TRP channel modulators as novel structures with potential therapeutic benefits.

## 1. Introduction

Hydrogen sulfide (H_2_S), an invisible gas with a rotten egg smell, is an occupational environmental toxicant (Rumbeiha et al. 2016). It is the 2^nd^ most common cause of fatal gas exposures in workplaces (Guidotti 2015). Recently, H_2_S gained attention for nefarious uses in chemical terrorism and in suicide (Binder et al. 2018; Morii et al. 2010). H_2_S is also endogenously produced at low concentrations in multiple organs, including the brain, serving multiple physiological functions in vertebrates. It is naturally synthesized from the amino acids, L-cysteine and L-cystathionine, catalyzed by multiple enzymes including cystathion-β-synthase, cystathionine-γ-lyase, and 3-mercaptopyruvate sulfur transferase (Polhemus and Lefer 2014).

Acute exposure to high concentrations of H_2_S induces severe neurotoxicity (Anantharam et al. 2018; Anantharam et al. 2017a; Anantharam et al. 2017b; Guidotti 2010; Kim et al. 2018; Kim et al. 2019; Rumbeiha et al. 2016; Snyder et al. 1995; Tvedt et al. 1991b). At concentrations higher than 500 ppm, H_2_S causes apnea, seizures, coma, and death (Guidotti 2015; Rumbeiha et al. 2016). Most deaths occur at the location of exposure (Santana Maldonado et al. 2022). Among survivors, acute H_2_S exposure often induces delayed neurological sequalae including nausea, persistent headache, movement disorders, impaired memory, amnesia, psychosis, anxiety, depression, sleeping disorders and coma. Some cases progress to prolonged vegetative states (Rumbeiha et al. 2016; Tvedt et al. 1991a; Wasch et al. 1989). Murine and porcine models of H_2_S poisoning exhibit seizure activity and loss of consciousness (Anantharam et al. 2018; Anantharam et al. 2017a; Anantharam et al. 2017b; Kim et al. 2018; Kim et al. 2019; O’Donoghue J 1961). Seizure activity has a high correlation with loss of consciousness in H_2_S-exposed mice (Anantharam et al. 2017a). The frequency and severity of seizure activity varies among mice, although seizure severity predicts shorter survival time (Anantharam et al. 2018; Anantharam et al. 2017a; Kim et al. 2018; Kim et al. 2019). Interestingly, preventing seizures with an anti-convulsant agent prolongs the survival of mice during H_2_S exposure (Anantharam et al. 2018).Victims of acute H_2_S poisoning also develop pulmonary edema (Tanaka et al. 1999) and heart failure (Sastre et al. 2013). Results from some animal models indicate cardiac arrest after acute sodium hydrosulfide (NaHS) exposure (Haouzi et al. 2015).

Despite this knowledge, it remains unclear what organ(s) primarily mediates H_2_S-induced physiological dysfunction leading to death and which are second order sequalae of H_2_S exposure. For example, respiratory arrest will eventually precipitate cardiac arrest as a secondary outcome, but the inverse is also true. Likewise, cardiac and respiratory arrest can both elicit cerebral hypoxia which can trigger electrocerebral suppression and, in extreme cases, seizures; however, seizures and hypoxia can also elicit cardiac and respiratory dysfunction. Determining which factors are initiating the H_2_S cascade necessitates simultaneous respiratory, cardiac and electrocerebral recordings under tightly controlled experimental conditions. Moreover, the molecular mechanisms underlying H_2_S-induced neurotoxicity and death are poorly understood.

It is widely reported that H_2_S inhibits cytochrome c oxidase enzyme in mitochondria, however, this alone does not explain the broad spectrum of toxic effects from knockdown and coma, to development of vegetative states (Anantharam et al. 2018; Anantharam et al. 2017a; Anantharam et al. 2017b; Guidotti 2015; Kim et al. 2018; Kim et al. 2019; Ng et al. 2019; Snyder et al. 1995; Tvedt et al. 1991a; Tvedt et al. 1991b). Transcriptomic and proteomic studies of the inferior colliculus in H_2_S-exposed mice showed that H_2_S induces dysregulation of multiple biological pathways including calcium homeostasis, oxidative stress, immune response, and neurotransmitters (Kim et al. 2018; Kim et al. 2019).

Calcium homeostasis is important for virtually all neuronal functions. Synchronous Ca^2+^ oscillations (SCOs) are ubiquitous signaling mechanisms whose frequency and amplitude encode information in both individual neurons (e.g., regulating physiologic metabolic and transcriptional responses) and at the level of networks (e.g., regulating synaptic connectivity and circadian rhythms) (Alford and Alpert 2014; Cavieres-Lepe and Ewer 2021; Tokumitsu and Sakagami 2022). SCO patterns encode frequency and amplitude signals that provide a means to control a wide range of cellular physiological functions (Smedler and Uhlen 2014; Sneyd et al. 2017). Dysregulation of SCO patterns has been reported in Alzheimer’s disease (Santos et al. 2009) and Huntington’s disease (Glaser et al. 2021). Various environmental toxicants have also been reported to disrupt SCO in neuronal cell cultures (Cao et al. 2014; Cao et al. 2017; Zheng et al. 2019).

Primary cortical neuronal/glial coculture (PCN) displays spontaneous electrical spike activity (ESA) that form synchronized and desynchronized field potentials as the PCN develops (Cao et al. 2014; Zheng et al. 2019). This PCN model exhibits spontaneous SCOs which are driven by ESA (Cao et al. 2014; Zheng et al. 2019) (Cao et al. 2012; Muramoto et al. 1993; Robinson et al. 1993). SCOs are shown to be strongly related to seizure activity in an in vitro epileptic model (Sombati and Delorenzo 1995). There are several types of Ca^2+^ channels in the brain that mediate physiological, pharmacological, and toxicological responses (Table 1). However, whether H_2_S alters SCO patterns and, if so, the channels contributing to dysfunction have not been investigated.

**Table 1.**
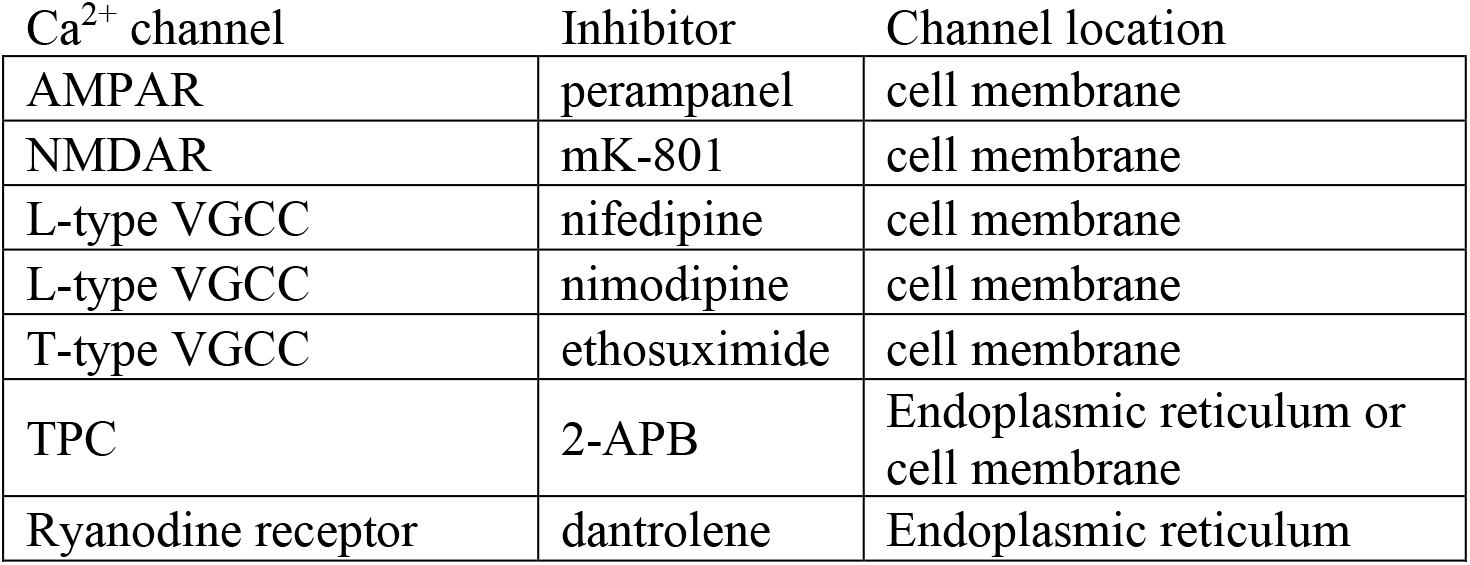
Select calcium channels in the brain, their specific inhibitors, and location

The present study uses an *in vivo* mouse model of inhaled H_2_S to measure brain (EEG), heart (EKG) and lung (plethysmography) to determine temporal changes during H_2_S exposure. We identify suppression of brain electrical activity (EEG) is the proximal effect of H_2_S intoxication preceding reduced pulmonary function and death. In combination with our prior knowledge that H_2_S causes Ca^2+^ dysregulation in brain and the important role of Ca^2+^ signaling in regulating neurological functions, we developed an in vitro PCN cell culture model to further test our *in vivo* findings of acute H_2_S exposure. The use of FLIPR and MEA assays with the PCN culture model provides new information about how H_2_S alters neuronal Ca^2+^ signaling and the mechanisms responsible.

## 2. Materials and Methods

### Reagents

Poly-L-Lysine (PLL), Fluo-4 AM, bovine serum albumin (BSA), cytosine β-D-arabinofuranoside (ARA-C), MK-801, nifedipine, nimodipine, 2-APB, and dantrolene were purchased from Sigma Aldrich (St. Louis, MO). Perampanel and ethosuximide were generously gifted from Dr. Michael Rogawski, University of California at Davis (UC Davis). Neurobasal media, HEPES, penicillin-streptomycin, B-27 Plus, GlutaMAX, fetal bovine serum (FBS), Flu-4 AM, calcein AM, and Hoechst 33342 were purchased from Thermo fisher scientific (Waltham, MA).

### Animals

Animal use was approved by the Institutional Animal Care and Use Committee (IACUC) of UC Davis or the University of Iowa, Iowa City. Animals were treated humanely and handled with care in accordance with IACUC guidelines.

### EEG, EMG and EKG surgery

C57BL/6J mice were purchased from Jackson Laboratories (000664; Bar Harbor, ME). Seven- to eight-week-old male C57BL/6J mice were housed at room temperature of 20-22 °C with a 12:12 h light dark cycle. Food (NIH-31 Mouse Diet) and water were available ad libitum. EEG and EKG electrodes were implanted as previously described (Muramoto et al. 1993). Briefly, mice were anesthetized under 1% isoflurane and skull surface was exposed and prepared. The headmount was then secured onto the skull aligning its middle with sagittal suture and the front two holes were in front of the coronal suture while the back two holes were in front of the lambdoid suture. Four pilot holes were bored with a 23G hypodermic needle. The screws were coated with silver epoxy and were driven into the holes acting as EEG electrode. Two stainless steel wires extended from the headmount were buried under cervical muscle to collect EMG signals. The headmount was then secured using dental cements. Mice were allowed to recover for at least 1 week before experimentation.

### EEG and EKG recording

EEG and EKG signals were acquired and processed as previously described (Purnell et al. 2017). Briefly, the preamplifier (8202-SL; Pinnacle Technology Inc.) was attached to the exposed plug on the implanted EEG headmount. EKG leads (MS303-76; Plastics One, Roanoke, VA) were implanted during the same surgery in the left chest wall and right axilla in a modified lead II configuration. The preamplifier cord was passed through an airtight gasket in the custom plethysmography chamber and then attached to a six-channel commutator (no. 8204; Pinnacle Technology Inc.). The signal was passed to a conditioning amplifier (model 440 Instrumentation Amplifier; Brownlee Precision, San Jose, CA). The EEG signals were amplified (100 times), band-pass filtered (0.3–200 Hz for EEG), and digitized (1,000 samples/s; NI USB-6008; National Instruments, Austin, TX). The digitized signal was transferred to a desktop computer and recorded using software custom written in MATLAB (R2018b; MathWorks, Natick, MA).

### Whole-body plethysmography

Whole-body plethysmography was employed to measure respiratory parameters during acute H_2_S exposure. Each animal was acclimated to the chamber. Baseline breathing was recorded for 10 min before exposure to 1000 ppm H_2_S or regular breathing air from pressurized cylinders. The recording chamber was outfitted with an ultralow volume pressure transducer (DC002NDR5; Honeywell International Inc., Morris Plains, NJ) to detect small pressure waves associated with breathing. The signal was amplified (100 times), band-pass filtered (0.3–30 Hz), and digitized before being recorded with a custom MATLAB software and stored on a desktop computer following previously published protocol (Purnell and Buchanan 2020). Before H_2_S or breathing air exposure, the signal was calibrated by delivering metered breaths (300 μl; 150 breaths min^−1^) via a mechanical ventilator (Mini-Vent; Harvard Apparatus) to the recording chamber. Respiratory function parameters including minute ventilation (VE), tidal volume (VT), and respiratory rate were assessed with MATLAB custom script as previously described (Hodges and Richerson 2008).

### Exposure paradigm

Mice were acclimated to the air-tight exposure chamber for one hour twice on different days prior to experimentation. EEG, EKG, and whole-body plethysmography recordings were calibrated before exposure to 1000 ppm H_2_S gas or breathing air from pressurized cylinders. Five minutes of baseline EEG, EKG, and whole-body plethysmography recordings were made before gas exposure. Gas exposure was simulated without an animal twice while the concentration of H_2_S in the plethysmography chamber was monitored in real-time using a H_2_S sensor (RKI instrument, Union city, CA). EEG, EKG, and breathing of C57BL/6J mice was monitored in real-time using implanted electrodes and the air-tight plethysmography chamber as shown in the exposure paradigm figure (Fig. 1A). Mice were continuously exposed to the gas until complete cardiac arrest as assessed by EKG (about 2 h) as the endpoint was mortality for this part of the study. Mice in the control group were exposed to normal breathing air, also from a pressurized cylinder.

**Fig. 1.**
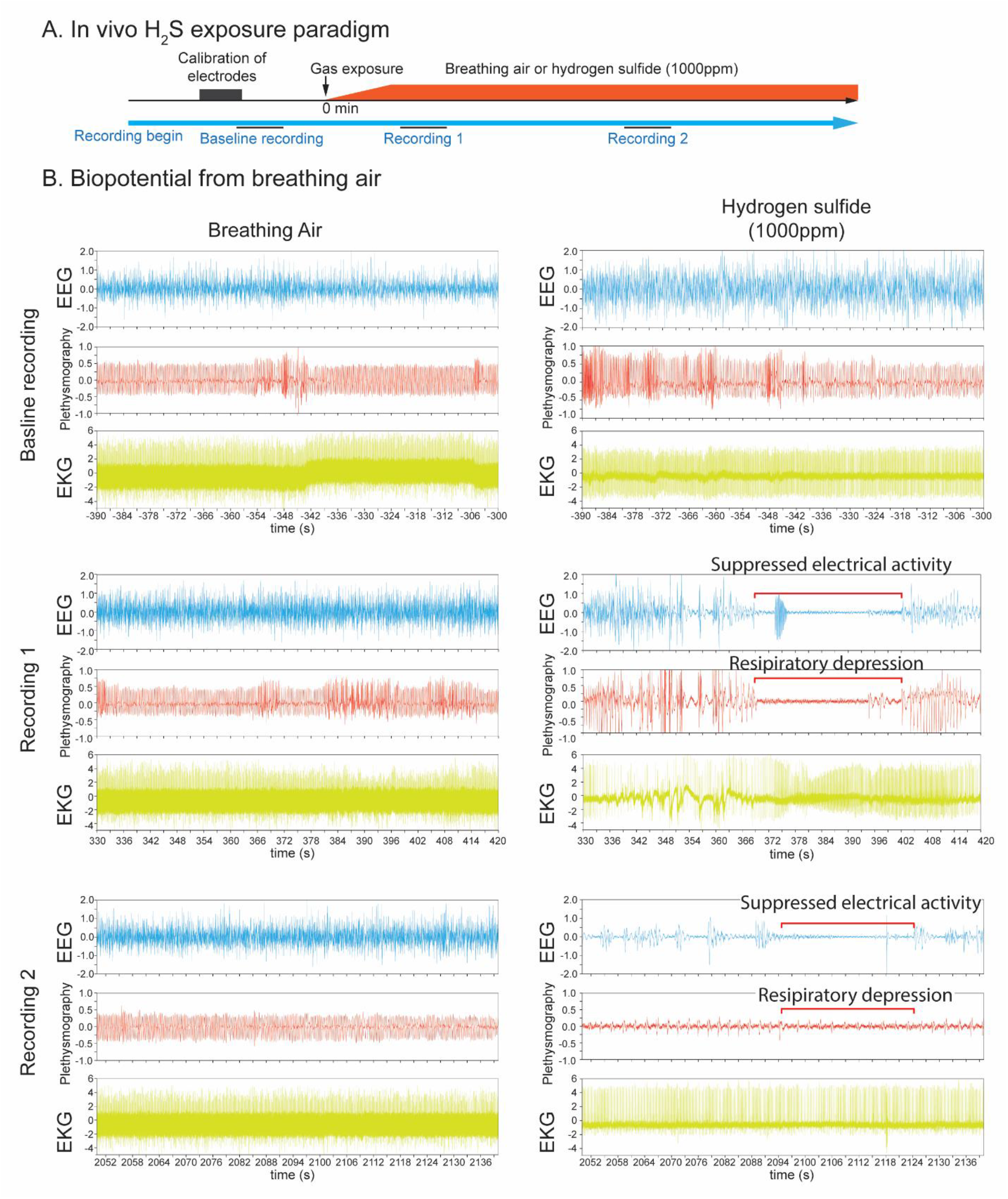
In vivo H_2_S-induced suppression of brain activity was concomitant with apnea. Mice were pre-implanted with electrodes for EEG and EKG. Breathing activity was recorded by plethysmography. In vivo H_2_S exposure paradigm was shown in Fig. 1A. After brief calibration of plethysmograph and recordings of biopotentials of baseline activity, mice were exposed to normal breathing air or 1000 ppm H_2_S (Fig. 1B) until death as assesses by EKG. Baseline recordings displayed EEG, EKG, and plethysmography traces of mice at rest. Recording 1 shows 90 sec signal recordings at about 5 min from the start of gas exposure, while Recording 2 shows 90 sec signal recordings at about 35 min from the start of gas exposure. Recordings 1 and 2 display typical suppression of EEG and plethysmography in H_2_S exposed mice (Fig. 1B). Note the H_2_S-induced electrocerebral suppression and respiratory arrest while cardiac activity persisted (Fig. 1B Recordings 1 and 2). N = 3 mice.

### Primary cortical neuron culture (PCN)

Male and female wild type (WT) C57BL/6J mice (000664) were purchased from Jackson Laboratories (Sacramento, CA) and were housed in the Teaching and Research Animal Care Services facility (TRACS) at the School of Veterinary Medicine, UC Davis with a 12:12 h light and dark cycle. Room temperature and relative humidity were maintained at 22 °C and 50 ± 10% respectively. Dissociated cortical neurons with minimal astrocyte composition were cultured as described previously (Cao et al. 2017). Briefly, neurons were dissociated from the cortex of C57BL/6J mouse pups of both sexes on postnatal day 0-1 and maintained in complete Neurobasal media [50 units of penicillin, 50 μg/ml of streptomycin, 2% (v/v) B-27 Plus, 10mM HEPES, and 1% (v/v) GlutaMAX] supplemented with 2% FBS. Dissociated cortical neurons were seeded onto the plate pre-coated with PLL at a density of 1 × 10^5^/well for 96 well plate for fluorescence laser plate reader (FLIPR Tetra; Molecular Devices, Sunnyvale, CA) or multi-electrode array (MEA) plate. The cell culture media was diluted to two-fold with complete Neurobasal medium the next day. A final concentration of 10 mM ARA-C was added to the culture medium at 36-48 h of culture to prevent astrocyte proliferation. The culture medium was changed every other day by replacing half volume of the culture medium with serum-free complete Neurobasal medium. PCN were maintained at 37°C with 5% CO_2_ and 95% humidity until experiment.

### Measurement of synchronous Ca^2+^ oscillations (SCOs)

Synchronous Ca^2+^ oscillations of the PCN culture were measured using fluorescent labeled calcium indicator and a FLIPR Tetra^®^ system as previously described (Cao et al. 2017). This fluorescent plate reader-based Ca^2+^ assay allowed measurement of functional Ca^2+^ signaling in PCN providing a rapid throughput in vitro system to assess the role of various Ca^2+^ channels in H_2_S-induced neurotoxicity using various pharmacological probes. Briefly, PCN between DIV 9 to DIV 16 were used. The growth medium was replaced with a dye loading solution (75 μL / well) containing 5 μM Fluo-4 AM and 0.5% BSA in Neurobasal medium. After incubation for 1 h in the dye loading solution, cells were washed 2 times with Neurobasal medium and once with imaging solution (125 mM NaCl, 5mM KCl, 2mM CaCl_2_, 1.2 mM MgCl_2_, 25mM HEPES, and 6 mM Glucose at pH 7.2). After washing, the cell medium was gently aspirated, and 125 μL of imaging solution was then added to the cells. The plate was transferred to FLIPR instrument to measure real-time intracellular Ca^2+^ concentration using a fluorescent signal. The fluorophore within the cells was excited at 488 nm, and Ca^2+^-bound Fluo-4 emission was recorded at 535nm. Baseline recordings of SCO were obtained before addition of the toxicant and/or drug probes using a programmable 96-channel robotic pipetting system. Properties of the SCO, including frequency and amplitude were analyzed using scipy module (version 1.7.1) in Python version 3.0 Software script (https://www.python.org/). A typical diagram of peak parameters of the normal SCO is shown in supplementary Fig. 1.

### Multi-electrode Array (MEA) recordings

Measuring spontaneous neuronal electrical activities using MEA system (Axion BioSystems, Atlanta, GA) was another cell function assay. It was performed as described previously (Cao et al. 2012). Briefly, dissociated PCN were seeded on to 48 well Maestro plates at a density of 1 × 10^5^ / well and maintained by changing growth medium every other day until use on 9-16 DIV. At the time of the experiment, the growth medium was gently aspirated and 125 μL Neurobasal medium was added to the cells. The plate was transferred to the MEA instrument pre-warmed to 37 °C. Cells were incubated on the MEA system for 5 min before measuring basal electrical activity. Cells were pre-treated with vehicle or pharmacological drug probes, followed by exposure to H_2_S. The MEA instrument recorded and amplified the raw extracellular electrical signals which were digitized in the Axis software at a rate of 25 kHz and filtered using a Butterworth band-pass filter (cutoff frequency of 300 Hz). The Axis software was used to detect spontaneous electrical events, perform spike analysis, and to export the analyzed data.

### Cell viability assay

Cell viability was performed using ImageXpress^®^ Micro (Molecular Devices, San Jose, CA) and cell-permeant dye, calcein AM, according to the manufacture’s protocol. Briefly, PCN were grown in pre-coated 96 well plate with PLL. Primary cortical neuronal/glial coculture DIV 9 to 16 were exposed to designated concentrations of H_2_S for 40 min at 37 °C. At the conclusion of the exposure period, the medium was aspirated and replaced with the growth medium. The H_2_S-exposed PCN were incubated for 72 h at 37 °C with change of growth medium after 48h. Calcein AM was applied to the H_2_S- exposed PCN and incubated for 30 min before taking images of cells using ImageXpress^®^ Micro. Hoechst 33342 was used for nuclei staining. Live cell counts were calculated using MetaXpress version 6 (Molecular Devices, San Jose, CA).

### Data analysis

Data are presented as mean and standard deviation of the mean. SCO peak parameters were normalized to the baseline values, which were compared to the control group. Statistical analyses of SCO peak parameters were performed by Student’s t-test using Microsoft Excel (Redmond, WA) or ANOVA using scipy statistical module (version 0.12.2) in Python version 3.0 (https://www.python.org/). A p-value of less than 0.05 was accepted as statistically significant.

## 3. Results

### 3.1 In vivo acute H_2_S exposure caused rapid suppression of electrocerebral and respiratory suppression which preceded cardiac dysfunction

Mice exposed to normal air showed normal patterns of EEG, EKG and plethysmography similar to the resting biopotentials throughout the observation period including recording epoch 1 and epoch 2 depicted in Fig. 1B. Exposure to 1000 ppm H_2_S induced significant disruption in brain and breathing activity starting within recording epoch 1 which became more severe within recording epoch 2 (Fig. 1B). Mice exposed to H_2_S exhibited an unstable burst-suppression pattern of EEG (Fig. 1B). Recording epoch 1 (Fig. 1B) shows that H_2_S-induced suppression of electrocerebral and respiratory suppression started as early as 5-7 min into H_2_S exposure. Thirty-four minutes into exposure (recording epoch 2), suppression of both electrocerebral activity and breathing became prominent (Fig. 1B). However, cardiac activity was substantially less strongly impacted. Mice exposed to H_2_S displayed frequent seizure activity which was followed by loss of postural control, profound electrocerebral suppression, and behavioral immobility, presumably due to a loss of consciousness. The duration of electrographic burst/suppression episodes varied, but typically lasted for a few seconds to about 30 sec. H_2_S exposed mice showed clear behavioral seizure activity before loss of consciousness characterized by myoclonus, body jerks, head bobbing, and loss of postural control; however, the cortical EEG traces during this time were not characterized by the high frequency, high amplitude discharges which typically characterized epileptiform discharges. The electrographic activity became progressively weaker as the exposure to H_2_S continued. Most notably, periods of electrographic suppression were frequently accompanied by apnea.

Respiratory suppression began early following the onset of H_2_S exposure and progressively worsened until the animal died (Fig. 2). In contrast, severe suppression of cardiac activity did not occur until > 30 min after the onset of H_2_S exposure (Figs. 2 and 3). Furthermore, the suppression of cardiac activity during the first 10 min of H_2_S exposure was partially recovered (Figs. 2 and 3). Given that respiratory suppression was only getting progressively more severe during this time period, it seems likely that the terminal decline in cardiac activity (> 30 min following exposure) was the consequence of prolonged hypoxia due to insufficient respiration (Fig. 2).

**Fig. 2.**
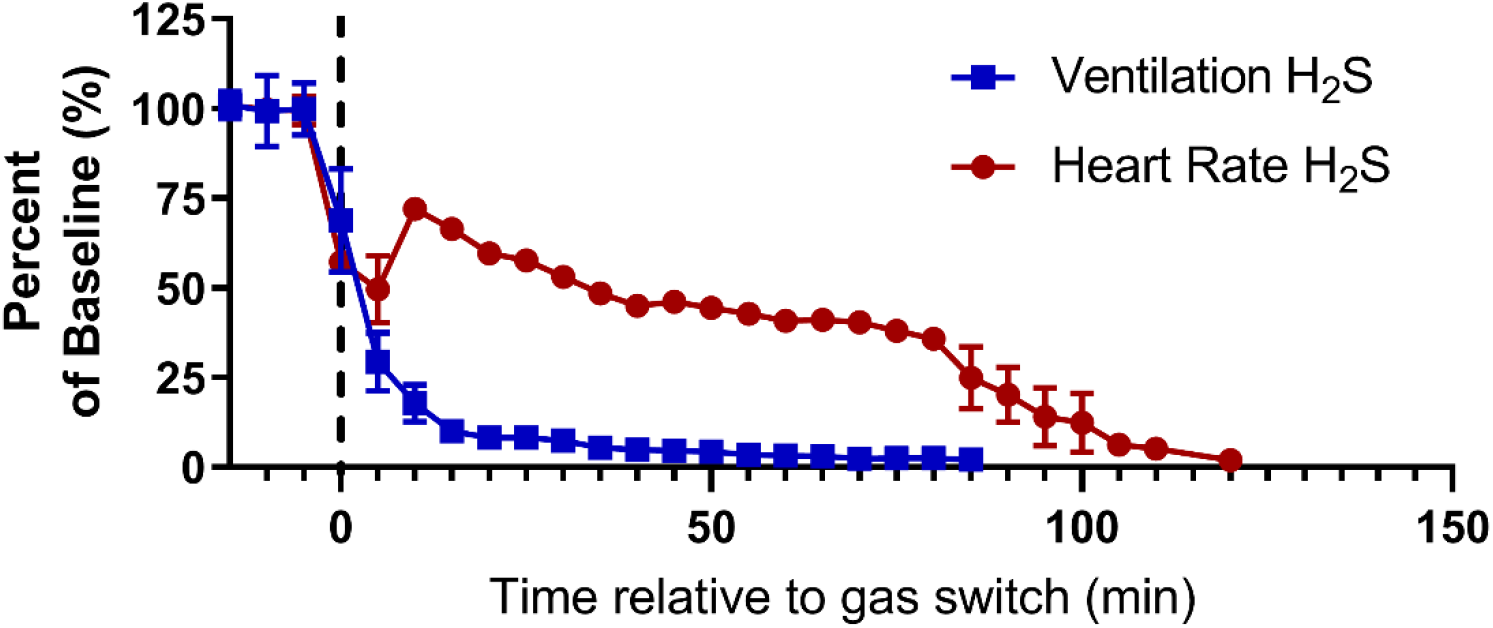
Respiratory suppression precedes cardiac suppression during H_2_S exposure. Time series data depicting baseline normalized ventilation (blue squares, mL/min/g) and heart rate (red circles; beats per minute) before and during exposure to 1000 ppm H_2_S. Data is depicted as mean with SEM, *n* = 3 mice. Baseline was defined as the 15 min prior to H_2_S administration. Grey dotted line, start of H_2_S exposure.

**Fig. 3.**
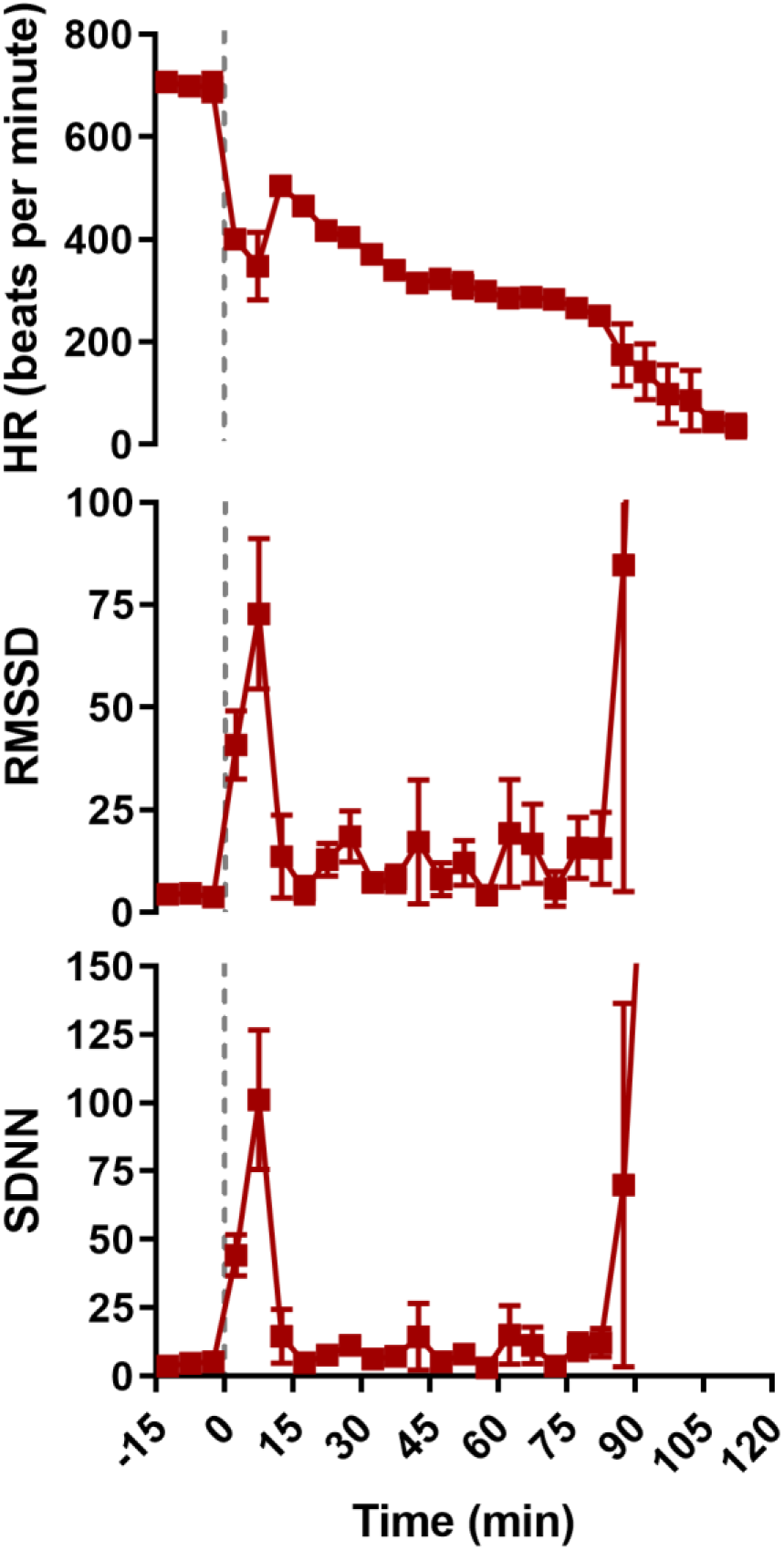
H_2_S exposure was characterized by an initial phase of decreased heart rate and increased heart rate variability. Time series data depicting heart rate (A; HR, beats per minute), root-mean square differences of successive R-R intervals (B; RMSSD, msec), and standard deviations of R-R intervals (C, SDNN, msec) before and during exposure to 1000 ppm H_2_S. Data was depicted as mean with SEM, *n* = 3 mice. Baseline was defined as the 15 min prior to H_2_S administration. Grey dotted line, start of H_2_S exposure.

### 3.2 H_2_S suppressed synchronous calcium oscillations (SCOs) in primary cortical neuronal networks (PCN)

SCOs in neurons play important roles in processing and interpreting sensory information in the central nervous system (CNS) (Muramoto et al. 1993; Robinson et al. 1993). The effects of H_2_S on SCO in PCN were examined using a rapid-throughput FLIPR Tetra^®^ system (Fig. 4). Cultured PCN DIV 9 to 16 were exposed to H_2_S from Na_2_S, a H_2_S chemical. Na_2_S is spontaneously converted to H_2_S in solution under physiological conditions (Equation 1).

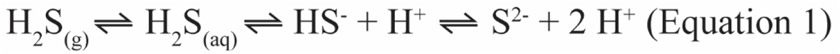

**Fig. 4.**
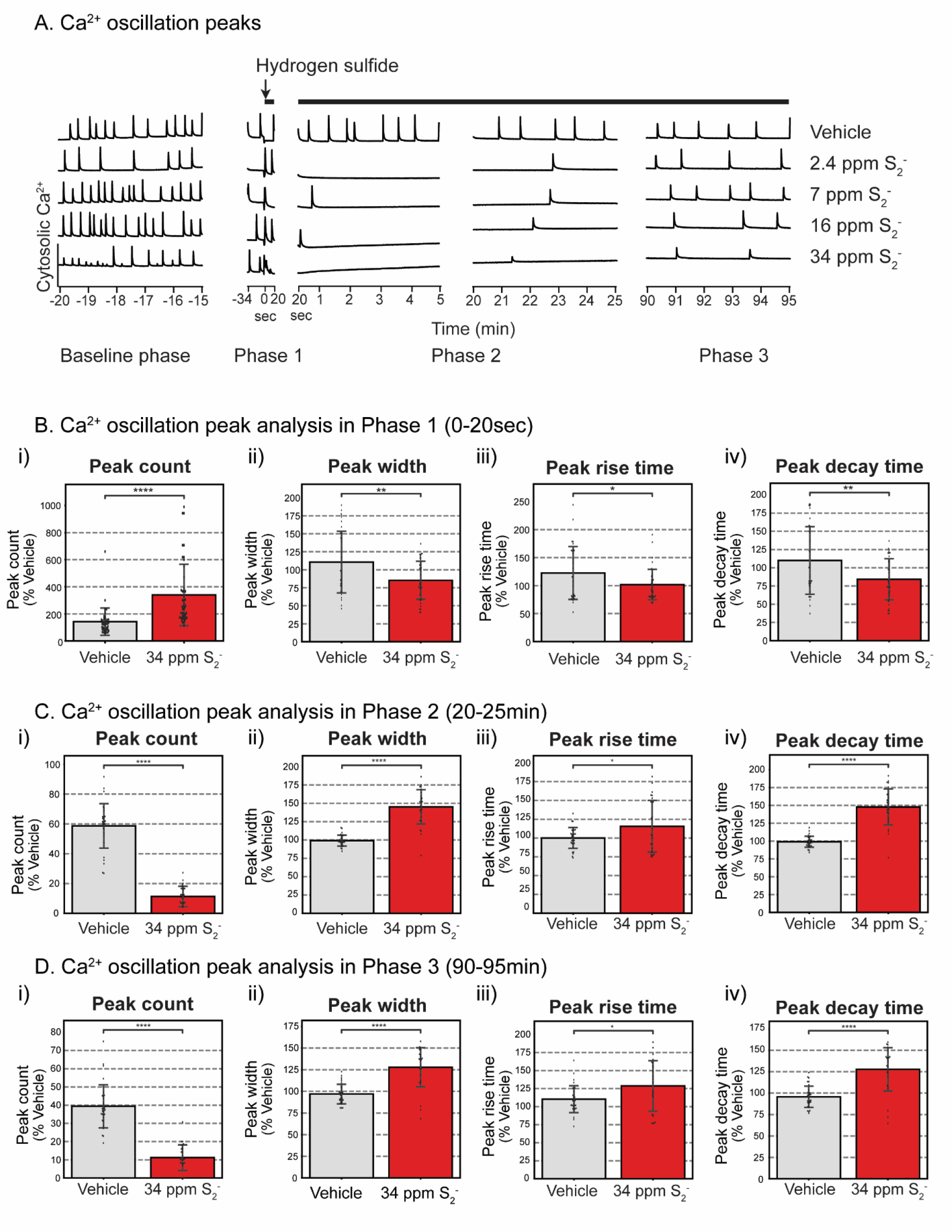
Synchronous calcium oscillations (SCO) in primary cortical neurons following exposure to H_2_S. Mouse PCN were exposed to graded concentrations of H_2_S. Intracellular Ca^2+^ levels were measured, and synchronous Ca^2+^ oscillations were analyzed. A) Traces of Ca^2+^ oscillations to show dose-response related effects of H_2_S exposure. A summary of SCO parameters in H_2_S exposed PCN for Phase 1 (0-20 sec), Phase 2 (20-25min), and Phase 3 (90-95min) is shown in graphs B, C, and D, respectively. Please note that H_2_S increased peak counts in Phase 1 but significantly suppressed this parameter in Phases 2 and 3. Also note that peak width, peak rise time, and peak decay time in Phase 1 were significantly reduced by H_2_S but significantly increased (reversed) in Phases 2 and 3. N > 30.

In pilot experiments, a broad concentration range of Na_2_S was prepared in imaging solution to identify the effective concentration-response relationship for use in our in vitro studies. In solution, H_2_S dissociates to hydrosulfide (HS^-^). At high pH (pH >12) HS^-^ is transformed to sulfide (S^2-^) ion. H_2_S generated from Na_2_S in the imaging solution was measured by adjusting the pH to a high pH (>12) to convert all HS^-^ to S^2-^ . We measured S^2-^ as a biomarker of H_2_S concentration in solution using a sulfide ion microelectrode. Changes in S^2-^ concentrations over the 60 mins for the dose-response study are summarized in Supplementary Fig. 2. Results show that the concentrations of S^2-^ decreased over the 60 min observation time, in a concentration dependent manner.

Effects of H_2_S by itself on SCO in PCN are summarized in Fig. 4. SCO activity at rest in PCN (Baseline phase) was measured 10 min before exposure to graded concentrations of H_2_S (Fig. 4A). During baseline, cultured PCN continuously exhibited baseline patterns of spontaneous SCO activity. Measured parameters included peak count, peak amplitude, peak width, peak rise time, and peak decay time. Exposure to vehicle (control) did not induce any significant changes in SCO peak parameters (Fig. 4A). However, addition of H_2_S induced a biphasic effect on the SCO in PCN. Initially, we observed an acute increase in SCO activity within the first 20-30 sec, followed by prolonged suppression of SCO (Fig. 4A). We named the acute hyper-active SCO phase as Phase 1, and the subsequent prolonged period of suppressed SCO as Phase 2 (Fig. 4A). We also noted that suppression of SCO started to recover at about 90 min after the addition of H_2_S. We called this recovery phase Phase 3 (Fig. 4A).

These effects of H_2_S were concentration dependent. Statistically significant differences between control and H_2_S exposed cells in each of the 3 phases are summarized in Figs. 4 B, C and D. Note that peak count was significantly increased by H_2_S in Phase 1 (Fig. 5Bi) but significantly suppressed in Phases 2 (Fig. 4Ci) and 3 (Fig. 4Di) compared to the respective Phases in vehicle control. H_2_S-exposed PCN showed significantly decreased SCO peak width, peak rise time, and peak decay time in Phase 1 (Fig. 4B ii, iii, and iv). These effects of H_2_S were reversed (increased) in peak width, peak rise time, and peak decay time in Phases 2 and 3 (Figs. 4C and D). Effects of H_2_S on SCO in Phase 2 were reminiscent of comma/depression phenotype in Recording 2 in Fig. 1C of the H_2_S exposed mice in vivo.

**Fig. 5.**
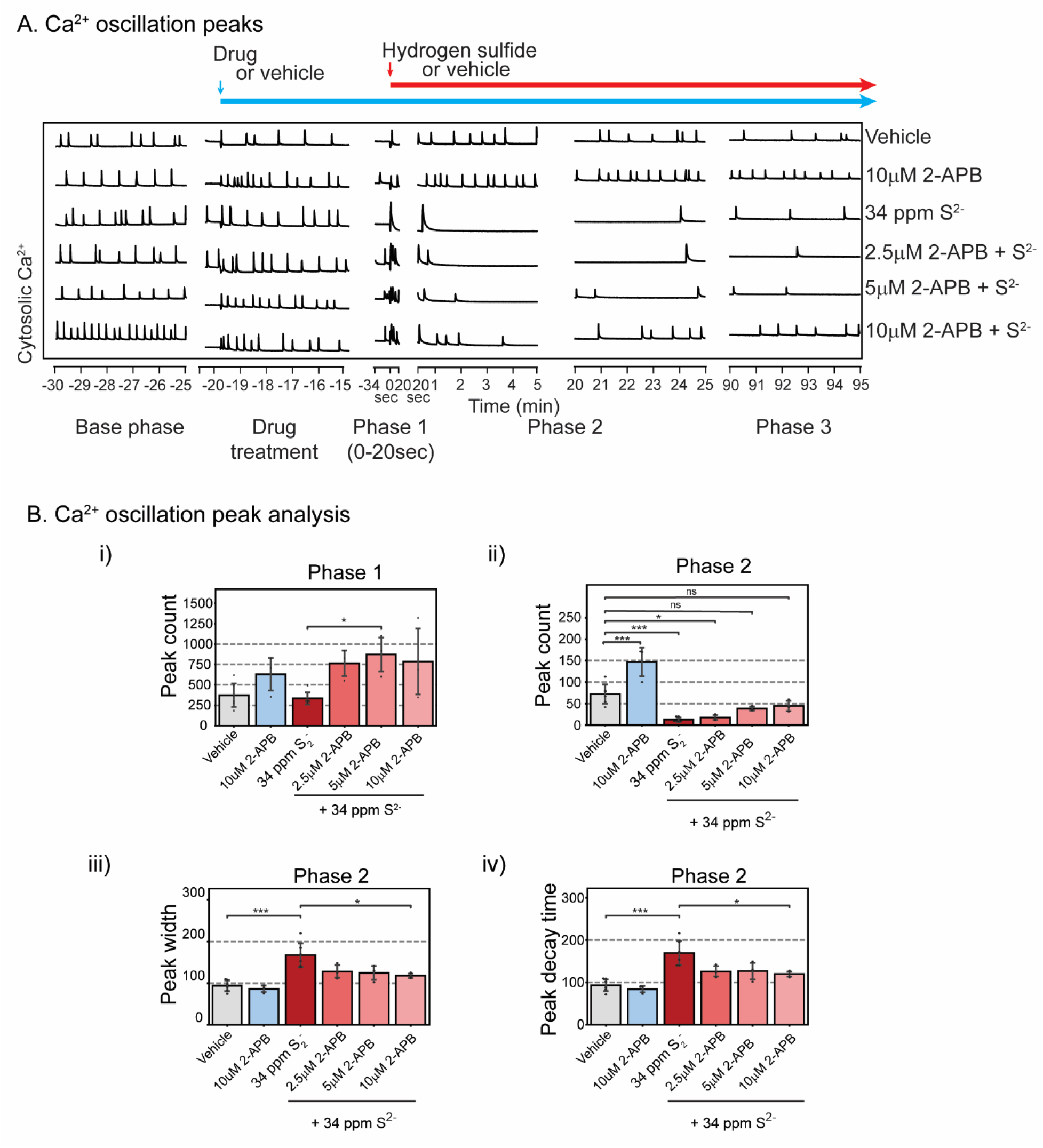
2-APB at 10 μM antagonized H_2_S-induced suppression of SCO in Phases 2 and 3 (A). Mouse PCN were pretreated with 2.5 - 10 μM 2-APB before exposure to H_2_S. Intracellular Ca^2+^ levels were measured, and synchronous Ca^2+^ oscillations were analyzed. 2-APB by itself at 10 μM significantly increased peak counts in Phase 2 (Bii). B) Analysis of SCO in H_2_S exposed cortical primary neurons. In Phase 2 2-APB at 10 μM significantly antagonized H_2_S-induced increase in peak width and peak decay time (Biii and Biv). Results in Phase 3 are similar to Phase 2 (data not shown). N = 3.

### 3.3 TRP channel blocker 2-APB prevents H_2_S-induced suppression of SCO

Store-operated Ca^2+^ (SOC) entry plays important roles in regulating intracellular Ca^2+^ in many eukaryotic cells, including neurons (Grudt et al. 1996). Of relevance to potential mechanisms contributing to high-level H_2_S toxicity observed in vivo is the emerging role of transient receptor protein (TRP) channels as primary mediators of physiological signals mediated by low levels of this gas in several organs, including the brain (Roa-Coria et al. 2019). We therefore examined whether 2-aminoethoxydiphenylborate (2-APB), a non-selective inhibitor of TRP channels, many of which function as store operated channels (SOC), (Baba et al. 2003) influence H_2_S-modified phases of SCO dysfunction described above. Mouse PCN were pretreated with 2.5 - 10 μM 2-APB before exposure to H_2_S mitigated the influences of H_2_S (Fig. 5A, representative traces). 2-APB by itself at 10 μM showed a tendency to increase peak count during Phase 1, although the difference did not reach statistical significance (Fig. 5Bi) until Phase 2 (Fig. 5Bii). 2-APB at 10 μM antagonized H_2_S-induced suppression of SCO in Phases 2 and 3. Intracellular Ca^2+^ levels were measured, and synchronous Ca^2+^ oscillations were analyzed. Pre-exposure to 10 μM 2-APB effectively antagonized the effects of H_2_S in Phases 2 and 3. Pretreatment with 2- APB dose-dependently increased peak counts in Phase 2 (Fig. 5Bii). Also in Phase 2, 2-APB at 10 μM significantly antagonized H_2_S-induced increase in peak width and peak decay time (Fig. 5Biii and 5Biv). Results in Phase 3 were like those in Phase 2 (data not shown).

To further investigate the role of endoplasmic reticulum (ER)-derived calcium in H_2_S-induced suppression of SCO we used dantrolene (DAN), a ryanodine receptor signaling inhibitor, as a pharmacological probe (Hayashi et al. 1997). Preliminary dose-range finding studies were done to optimize the concertation of DAN for use in this experiment. It is interesting that DAN did not show efficacy in preventing H_2_S-induced suppression of SCO (Fig. 6A).

**Fig. 6.**
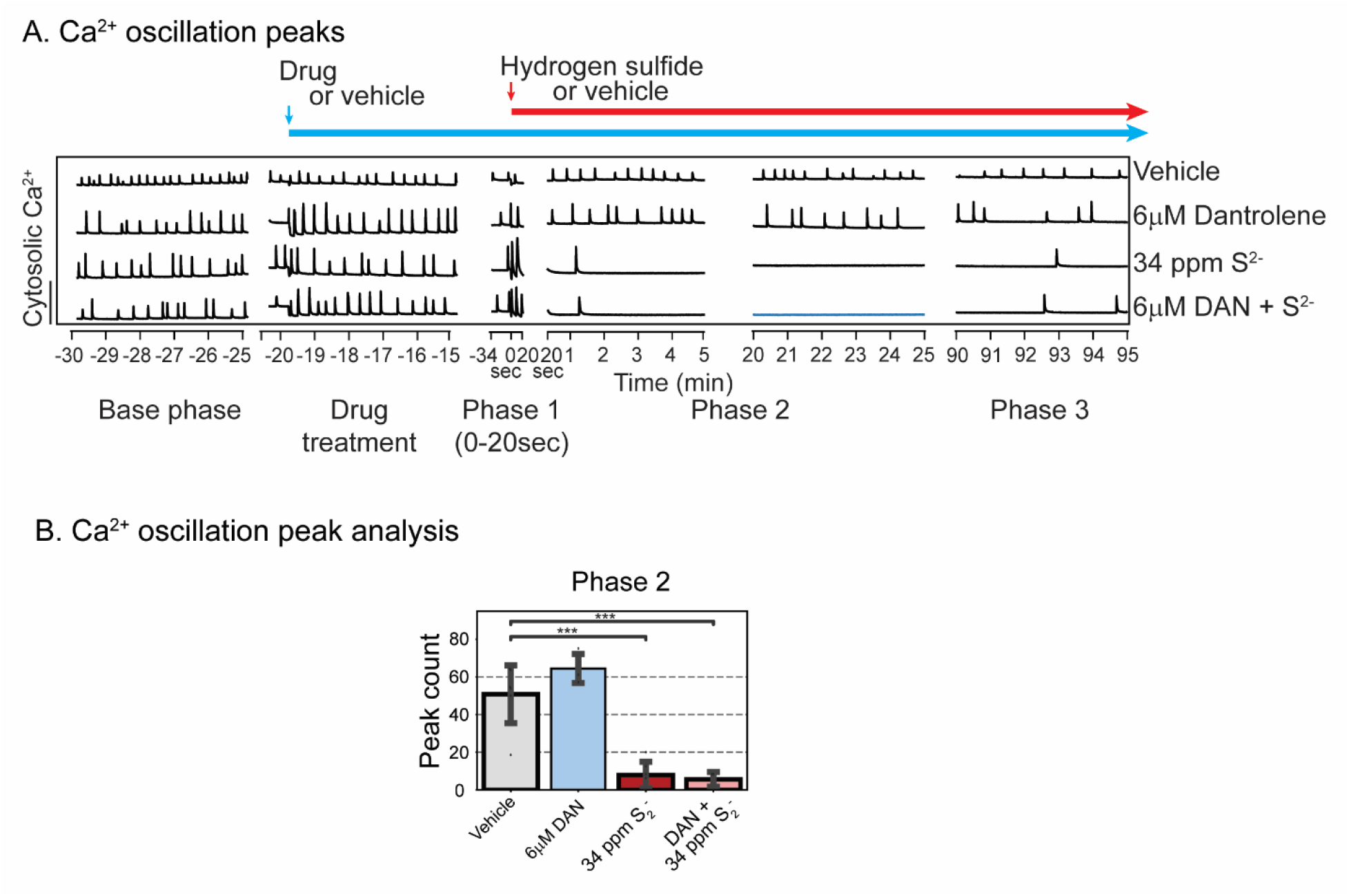
Mouse PCN were pre-treated with 6 μM dantrolene (DAN) before exposure to H_2_S. Intracellular Ca^2+^ levels were measured, and synchronous Ca^2+^ oscillations were analyzed. A) Traces of Ca^2+^ oscillations are shown. Note that pre-treatment with Dantrolene had no impact on H_2_S-induced SCO in Phases 2 and 3. B) A summary of SCO peak counts in H_2_S exposed cortical primary neurons in Phase 2. Note that DAN failed to prevent on H_2_S-induced suppression of peak counts. Similar results were observed for peak width and peak decay time in Phases 2 and 3 (data not shown). N = 3.

### 3.4 Ca^2+^ channel blockers differentially influence abnormal SCO patterns triggered by H_2_S

We proceeded to investigate pharmacological blockers with a spectrum of selectivity towards a number of neuronal voltage-dependent Ca^2+^ channels to determine whether the mitigating effects observed with 2-ABP could be generalized. Basal SCO activities were recorded prior to application of pharmacological drugs as depicted with baseline shown in Fig. 7A. PCN were pre-treated with nifedipine (Nif, 1-10 μM), followed by exposure to H_2_S (34 ppm S^2-^). SCOs in each phase were assessed (Fig. 7B and C). Pre-treatment with Nif significantly and dose-dependently increased acute H_2_S-induced SCO peak counts in Phase 1 compared to vehicle (control) group (Fig. 7Bi). Although Nif dose-dependently reduced peak width and peak decay time in Phase 1, these were not statistically significant compared to vehicle control (Fig. 7B ii and iii).

**Fig. 7.**
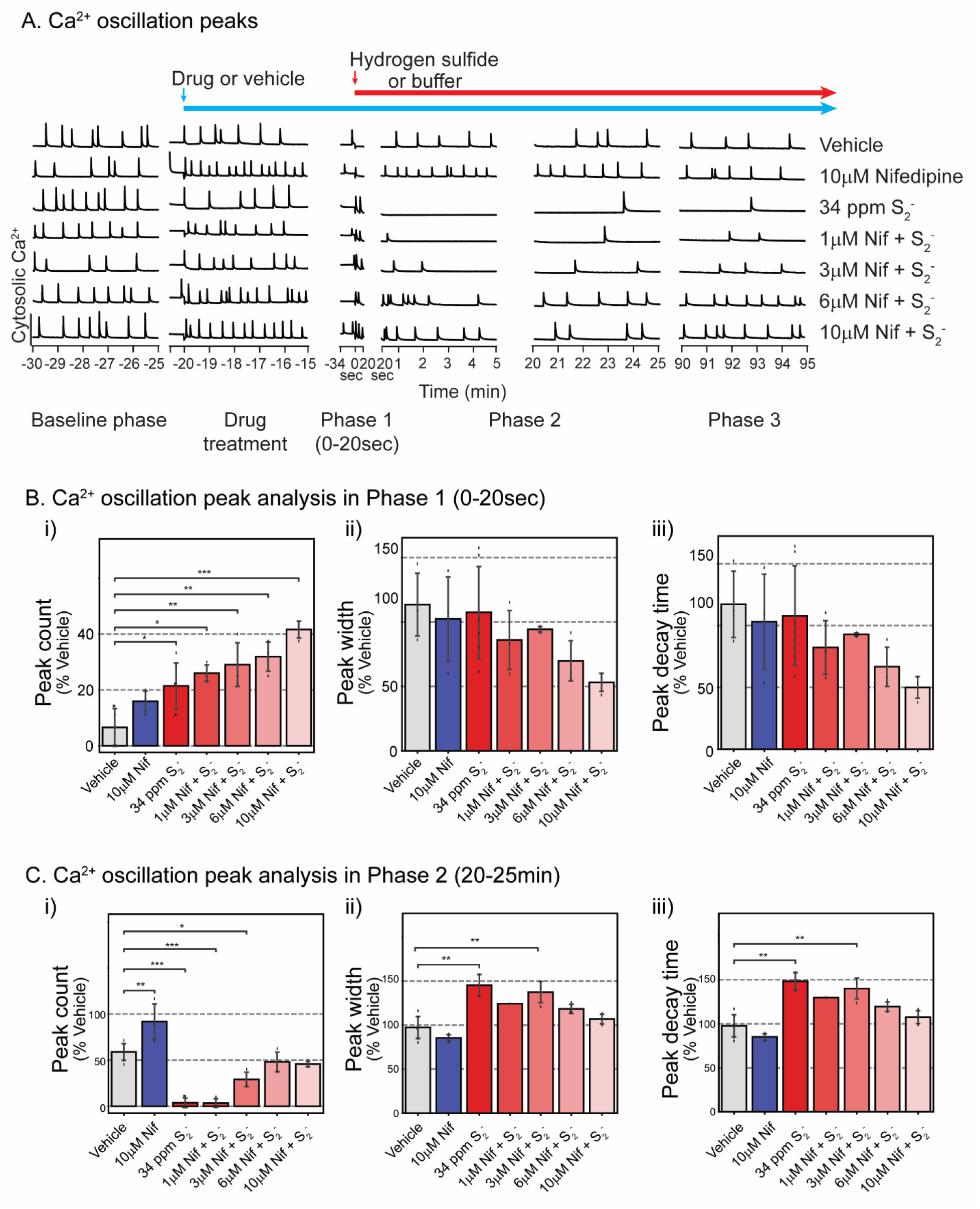
Pre-exposure to nifedipine (Nif), an inhibitor of L-type voltage dependent calcium channel, antagonized H_2_S-induced suppression of SCO in Phases 1-3 at 6 and 10 μM in mouse PCN. Intracellular Ca^2+^ levels were measured, and SCO were analyzed. A) Traces of Ca^2+^ oscillations showing that Nif at 6 and 10 μM effectively prevented H_2_S-induced dysregulation of SCO. A summary of SCO characteristics in H_2_S exposed PCN for Phase 1 (0-20 sec) and Phase 2 (20-25min) in B and C. Effects of Nif in Phase 3 were similar to those in Phase 2 (data not shown). N = 3.

Consistent with results shown in Fig. 4 H_2_S suppressed peak counts in Phase 2 but pretreatment with Nif antagonized this H_2_S-induced effect at 6 and 10 μM concentrations (Fig. 7Ci). In Phase 2 H_2_S increased both peak width and peak decay time consistent with observations in Fig. 4 Notably, pretreatment with 6 or 10 μM Nif prevented H_2_S-induced increase in peak width (Fig. 7C ii) and peak decay time (Fig. 7C iii). Effects of nifedipine in Phase 3 were like those in Phase 2 (data not shown). An interesting observation was that pretreatment with nimodipine, a chemically related L type voltage-gated Ca^2+^ channel inhibitor, failed to prevent H_2_S-induced suppression of SCO in PCN (Fig. 8Aiii and 8Biii).

**Fig. 8.**
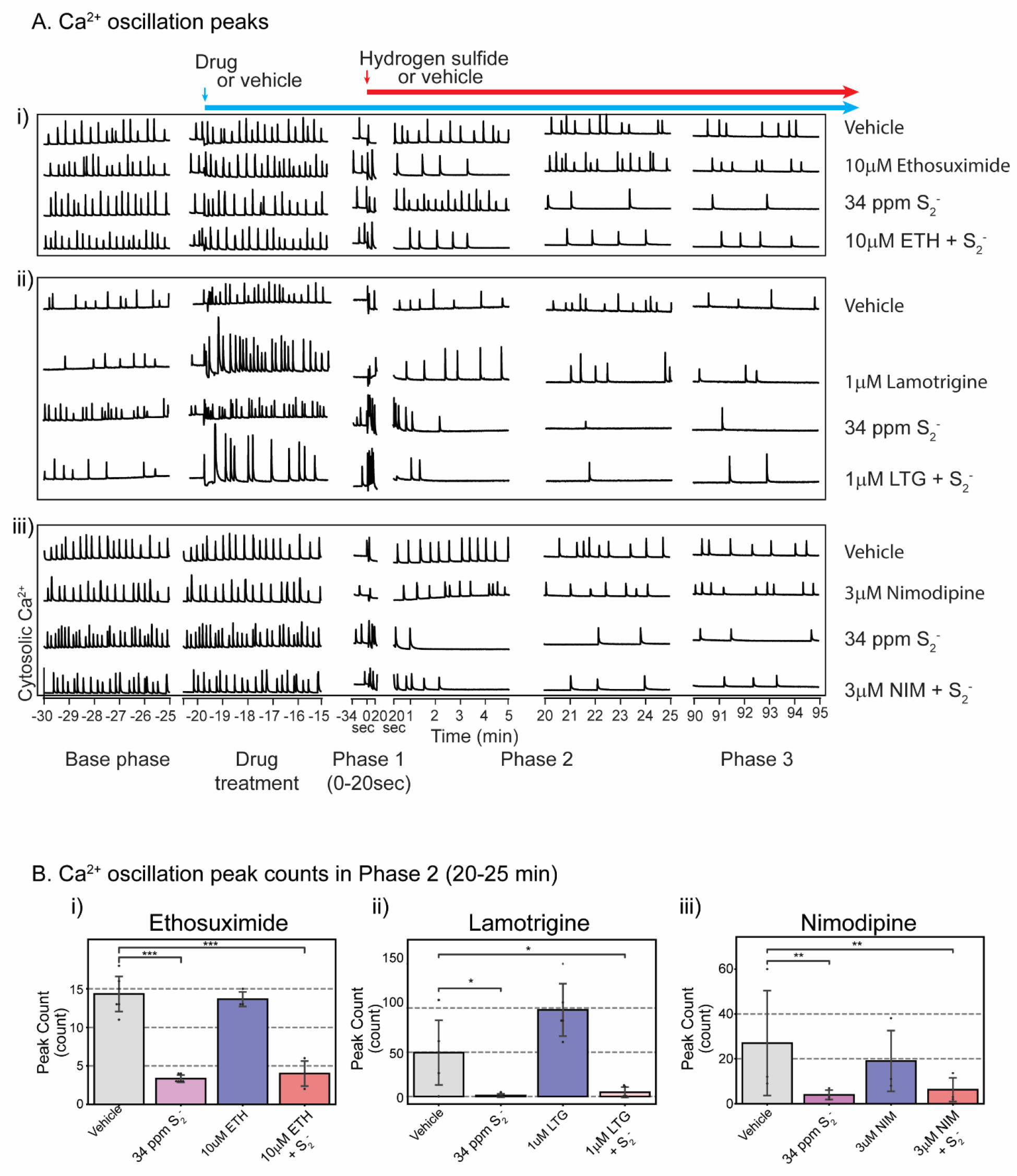
Ethosuximide, lamotrigine, and nimodipine failed to prevent H_2_S-induced suppression of SCO. Mouse cortical primary neurons were pre-treated with ethosuximide, lamotrigine, or nimodipine before exposure to H_2_S. Intracellular Ca^2+^ levels were measured, and synchronous Ca^2+^ oscillations were analyzed. A) Traces of Ca^2+^ oscillations. B) A summary of analyses of SCO activity in H_2_S exposed cortical primary neurons following pretreatment with ethosuximide, lamotrigine, and nimodipine. N = 3.

To investigate the role T type Ca^2+^ channels play in H_2_S-induced neurotoxicity, ethosuximide (ETH), a T type Ca^2+^ channel blocker, was used. As was the case for Nif and nimodipine, preliminary dose-range finding studies were done to select the ideal test dose for ethosuximide, lamotrigine and nimodipine. Pre-treatment with ethosuximide did not prevent H_2_S-induced suppression of SCO peak counts (Fig. 8Ai and 8Bi). We then used lamotrigine to investigate the effects of sodium channels on H_2_S-induced toxicity. Lamotrigine (LTG) is a voltage-dependent sodium channel blocker (Cheung et al. 1992). Pre-treatment with lamotrigine also failed to antagonize the effects of H_2_S-induced dysregulation on SCO in both Phase 1 and Phase 2 (Fig. 8Aii).

### 3.5 NMDA and AMPA receptor antagonists failed to prevent H_2_S-induced suppression of SCO

It has been reported that H_2_S toxicity is mediated by glutamate (Cheung et al. 2007) and that it induces influx of Ca^2+^ through NMDA receptors. MK-801, an NMDA receptor inhibitor, was used to investigate the role of NMDA receptors in H_2_S-induced suppression of SCO (Fig. 9). As with other pharmacological probes, preliminary range finding studies were performed to select the optimal concentration for MK-801 and perampanel (PRP; an AMPA receptor blocker). (Joshi et al. 2011) Pre-treatment of PCN with MK-801 alone induced suppression of SCO. Addition of H_2_S exacerbated the MK-801-induced suppression of SCO in PCN (Fig. 9A i and B ii). Pre-treatment of PCN with PRP, suppressed SCO activity (Fig. 9A ii and 9B iii and iv). Unique among all drug probes used in this study, PRP prevented H_2_S-induced increase in peak counts in Phase 1 (Fig. 9B iii). However, in Phase 2 pre-treatment with PRP augmented H_2_S-induced suppression of SCO in PCN (Fig. 9B iv).

**Fig. 9.**
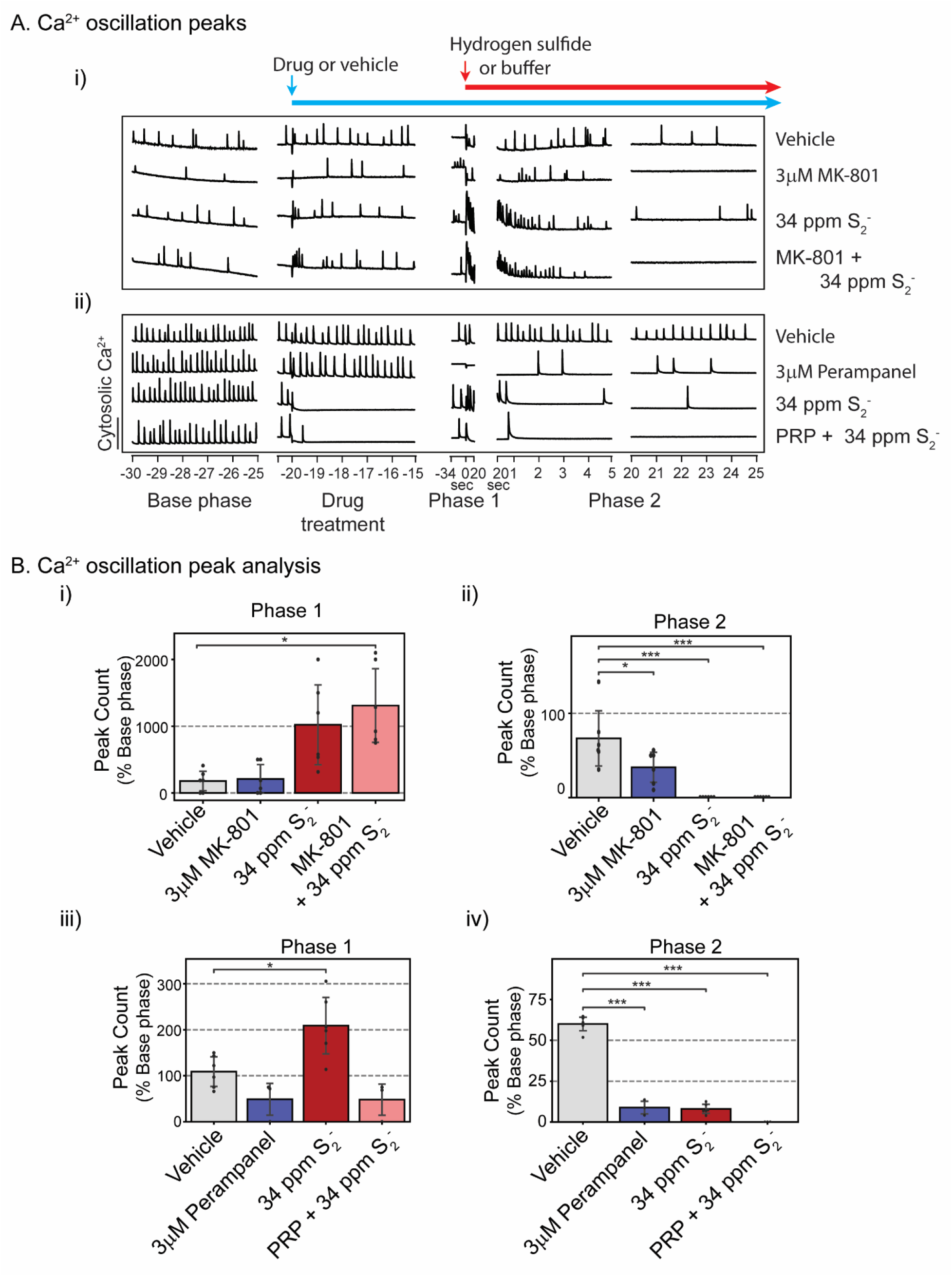
Mouse PCN were pretreated with MK-801 or perampanel (PRP) before exposure to H_2_S. Intracellular Ca^2+^ levels were measured, and synchronous Ca^2+^ oscillations were analyzed. A) Traces of Ca^2+^ oscillations were shown. Inhibitors for NMDA receptor and AMPA receptor failed to antagonize H_2_S -induced suppression of SCO in Phase 2 (Fig. A). B) Analysis of SCO in H_2_S exposed PCN. MK-801 failed to prevent H_2_S-induced effects on peak counts in Phases 1 and 2 (Figs. Bi and Bii). PRP significantly prevented H_2_S-induced increase in peak counts in Phase 1 (Fig. Biii) but worsened H_2_S-induced decrease in peak counts in Phase 2 (Fig. Biv). N = 3.

### 3.6 Pretreatment with 2-APB prevented H_2_S-induced suppression of neuronal electrical network spike activity in primary cortical neurons using MEA

Considering the overall activity of 2-APB towards mitigating the three phases of H_2_S-triggered SCO dysfunction, we plated PCN on multi-electrode arrays (MEA) to measure electrical spike activity (ESA) (Johnstone et al. 2010). Spontaneous neuronal ESA in PCN at rest was measured before treatments to obtain 10 min of baseline. Following recording of baseline activity, PCN were pretreated with 10 μM 2-APB or buffer 20 min before addition of H_2_S (34 ppm S^2-^). Recording of neuronal electrical activity was initiated 1 min before H_2_S was added (Fig. 10A). Neuronal spike activity was detected by the Axis software. Addition of H_2_S immediately induced hyperactive electrical stimulation by more than 75% compared to vehicle group (Fig. 10A and Bi) which was followed by suppression of electrical activity (Fig. 10A and Bii). We named the hyperexcitation period as Phase 1 and the suppression phase as Phase 2, similar to our data analysis strategy with FLIPR. Pretreatment with 2-APB prevented H_2_S-induced acute hyper-active electrical activity in Phase 1. Whereas H_2_S significantly suppressed neuronal electrical activity in Phase 2 (20 min after H_2_S exposure), 2-APB pre-treatment modestly prevented H_2_S-induced suppression of neuronal electric activity by 27% compared to the vehicle group (Fig. 10B ii).

**Fig. 10.**
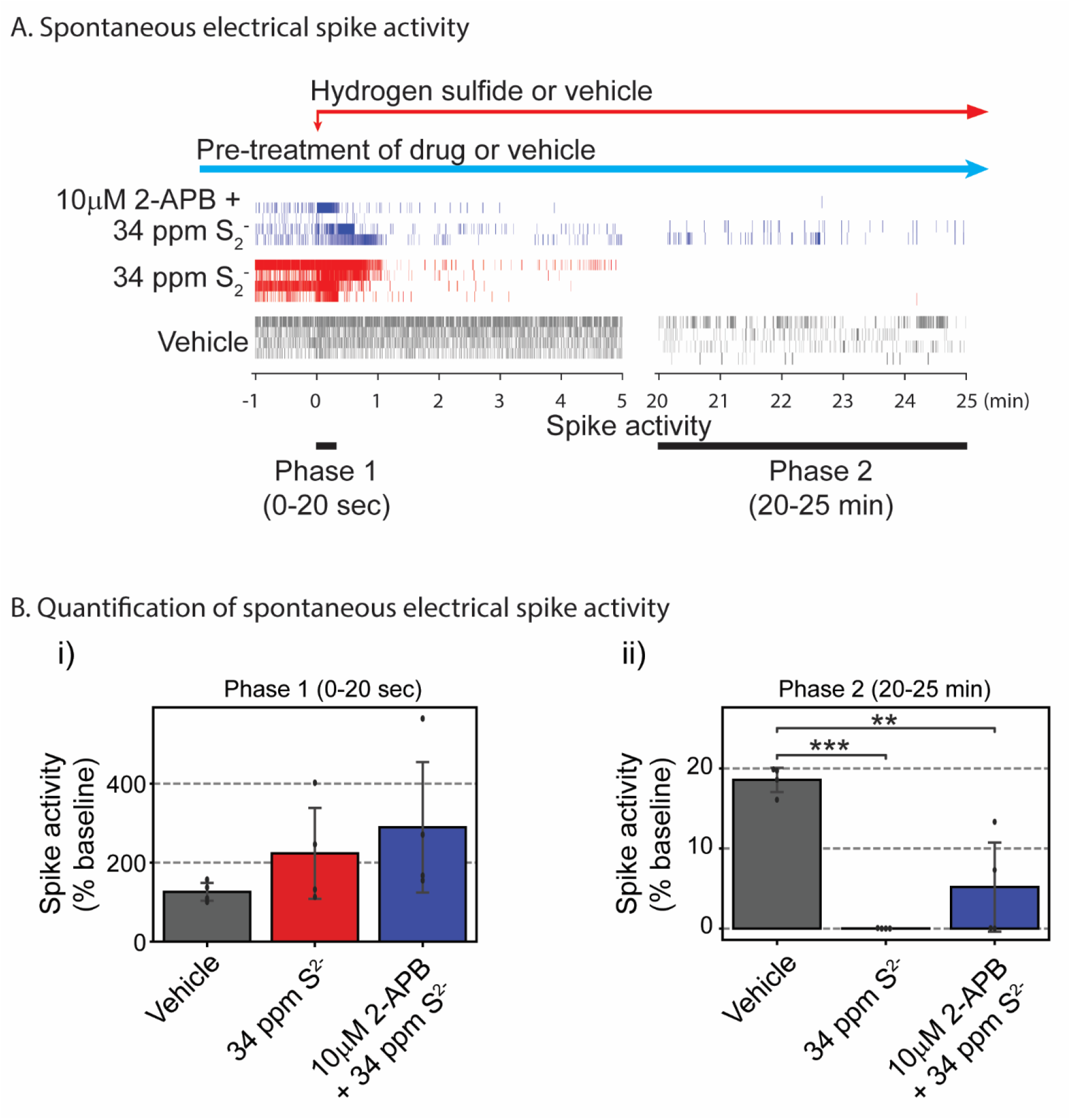
Neuronal electrical activities were measured using the multiple-electrode array system (MEA). Mouse PCN were pre-treated with 2-APB or vehicle before exposure to a H_2_S donor. A) spontaneous neuronal electrical spike activities are shown. Addition of H_2_S induced hyperactive electrical spike activity in Phase 1, whereas pretreatment with 2-APB modestly antagonized H_2_S-induced suppression of electrical activity in Phases 1 and 2. B) Spontaneous electrical spike activities in PCN were quantified. Note that pretreatment with 2-APB modestly antagonized H_2_S-induced suppression of electrical activity in Phase 2 (Fig. 5Bii). N = 4.

### 3.7 2-APB protected cortical neurons from H_2_S-induced cell death

Of the diverse ion channel probes investigated in this study, 2-APB was the most interesting as far as antagonizing effects of H_2_S on SCO. We therefore investigated whether the antagonistic effects of 2-APB observed in this study had impact on H_2_S-induced neuronal cell death. To do this, a cell viability assay was performed using ImageXpress^®^ Micro (Molecular Devices, San Jose, CA) and cell-permeant dye, calcein AM, according to the manufacture’s protocol. (Fig. 11). PCN DIV 9-15 were pre-treated with 2-APB or buffer 10 min before exposure to H_2_S (34 ppm S^2-^) for 40min. The H_2_S exposure medium was then replaced with fresh cell growing medium. Cells were incubated for 72h after H_2_S exposure before assessing cell death by counting the live cells using fluorescent calcein dye (Fig. 11A). A summary of results of live cells is shown in Fig. 11B. H_2_S by itself induced about 50% cell mortality. For cells pretreated with 10 μM 2-APB, only 25% cell mortality was observed (Fig. 11B). These preliminary results suggest that 2-APB was neuroprotective in this model.

**Fig. 11.**
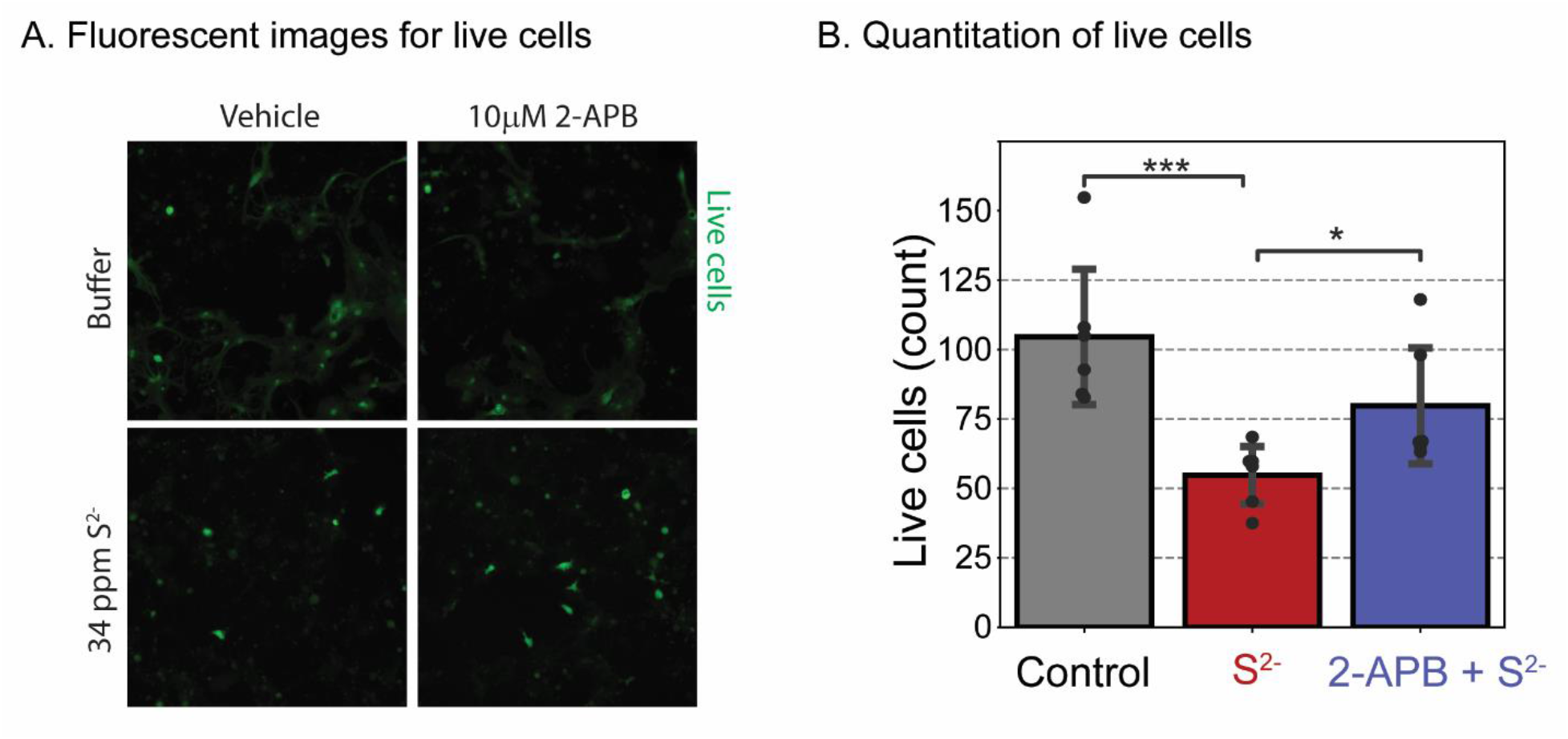
Mouse PCN were pre-treated with 10 μM 2-APB and exposed to H_2_S for 40 min only. The cells were washed and incubated for 72h before imaging of live cells to assess cell survival. A) Live cortical neurons were imaged using cell-permeable calcein AM fluorescent dye. B) A summary of results. Pretreatment with 2-APB significantly increased cell survival compared to the H_2_S group. N = 6.

## Discussion

The toxicity of H_2_S is complex and the proximate cause of death following acute H_2_S exposure is debatable. Moreover, the molecular mechanisms of H_2_S-induced neurotoxicity are not known. The study addressed several of these gaps in knowledge. First, in the in vivo study we show that it is electrocerebral suppression which precipitates respiratory suppression and outright apnea. Cardiac arrest is of later onset and likely a consequence of hypoxia due to prolonged respiratory insufficiency and/or H_2_S-induced lack of ATP. The onset of H_2_S exposure mildly decreased cardiac activity and increased heart rate variability. These mild changes in heart function partially recovered early during exposure. In contrast, respiratory suppression was severe and did not recover (Figs. 2 and 3). H_2_S exposure rapidly elicited a burst-suppressed EEG pattern in which periods of electrographic suppression frequently coincided with apnea (Fig. 1). As the exposure progressed, periods of bursting became less regular giving way to complete electrocerebral suppression (Fig. 1). This progressive electrocerebral dysfunction was temporally mirrored by progressive decreases in breathing until respiration was no longer compatible with life (Figs. 1 and 3). It is interesting however that we did not see clear epileptiform discharges in this study, suggesting that the electrographic seizure activity was localized to the brainstem. This is supported by our observations that seizure activity in this model are akin to audiogenic induced seizures in DBA mice and other models which originate from the brainstem.

Respiratory rhythmogenesis is a function of the brain, specifically pre-Bötzinger complex within the ventrolateral medulla (Smith et al. 1991). The coincidence between episodes of electrocerebral suppression and frank apnea may be indicative of transient periods neuronal inactivity across the entirety of the brain, including the respiratory nuclei of the brainstem. The data presented here indicate that (1) the electrocerebral dysfunction induced by H_2_S exposure is the driving force behind the respiratory dysfunction and (2) the cardiac dysfunction observed in the later stages of H_2_S exposure are causally downstream to the electrocerebral/respiratory dysfunction.

To further understand the mechanisms involved in electrocerebral suppression, we developed an in vitro model of PCN which recapitulated the in vivo H_2_S-induced EEG using SCO in FLIPR and electrical spike activity using MEA. In this study we used PCN because primary neuron/astrocyte coculture have functional receptors, unlike immortalized cell lines. FLIPR is a fluorescent plate reader-based assays that allows fast and simultaneous readings of intracellular Ca^2+^ of cells in multiple wells. Use of PCN with rapid-throughput systems using FLIPR and MEA allowed us to gain insights in the role of the various calcium channels in acute H_2_S- induced toxicity on the brain.

Synchronous calcium oscillation (SCO) is driven by action potentials in the neurons. The infinite patterns of frequency of SCO and changes of SCO frequency or amplitude provides important means to fulfil diverse cellular and tissue functions (Smedler and Uhlen 2014). Dysregulation of SCO in neurons is also implicated in toxicant-induced neuronal dysfunction (Cao et al. 2012). As shown in FLIPR experiments, H_2_S induced an acute hyper-active neuronal firing upon addition of H_2_S (Phase 1), followed by suppression of spontaneous neuronal firings in MEA system (Phase2). The acute hyper-active SCO and neuronal firing is akin to the seizure like burst suppression activity that was observed in vivo whereas suppressed SCO and suppressed neuronal firings are akin to loss of consciousness or central nervous system depression characteristic of acute H_2_S poisoning.

Acute hyper-active SCO in Phase 1 was characterized by increased SCO peak count and a shorter SCO peak rise time, width, and decay time indicating the cycle of SCO is much faster, which is also corresponds with the acute hyper-active neuronal firings in the MEA system. Phase 2 SCO is characterized by wider peak width and by increased SCO peak decay time. The finding that the SCO peak rise time in Phase 2 is not significantly altered while SCO peak decay time is increased indicates that H_2_S may affect the release intracellular calcium. Action potential drives SCO in cortical neurons (Robinson et al. 1993), and action potential initiates opening of sodium channels in neurons (Grider et al. 2022). We hypothesize that the acute hyper-active SCO plays a role in prolonged suppression of SCO in PCN. Treatment with 1 μM lamotrigine, a sodium channel blocker, by itself significantly decreased SCO peak count in Phase 1 while 30 μM lamotrigine completely abolished SCO in Phase 1 (data not shown). However, lamotrigine failed to prevent H_2_S-induced acute hyper-active SCO in Phase 1, within 20 sec of adding H_2_S (data not shown). These results indicate that H_2_S may induce acute hyper excitability in neurons in Phase 1 via a different mechanism other than through the lamotrigine-sensitive sodium channel.

Neurons have numerous Ca^2+^ channels targeted to specific anatomical regions that mediate specific physiological functions (Berridge 2016; Brini et al. 2014). We investigated select Ca^2+^ channel blockers for their ability to reverse H_2_S-induced dysregulation of SCO including acute hyper-active (Phase 1), prolonged suppression (Phase 2) and recovery (Phase 3) SCO epochs that we identified as biomarkers of cellular dysfunction using the FLIPR Tetra^®^ imaging system. We observed that not all Ca^2+^ channel blockers had the same influence on the three phases of H_2_S-triggered Ca^2+^ dysregulation. Moreover, those that showed efficacy toward antagonizing effects of H_2_S did not reverse all 3 phases of dysfunction. For example, nifedipine failed to suppress acute hyper-active SCO in Phase 1 but it reversed H_2_S-induced suppression of SCO in Phase 2. It is not clear what the exact mechanism(s) behind nifedipine-induced reversal of suppressed SCO by H_2_S exposure are at this time. Further research is needed to understand more about nifedipine-induced reversal of SCO when PCN culture was exposed to H_2_S. Although both nifedipine and nimodipine are L type VGCC inhibitors and are used for anti-seizure treatment, they showed different efficacy towards seizures(Grabowski and Johansson 1985; Konrad-Dalhoff et al. 1991). Nimodipine has a higher efficacy towards blocking Ca_V_1.3_α1_ (Xu and Lipscombe 2001), whereas nifedipine blocks Ca_V_ 1.2 at lower concentrations that nimodipine (Wang et al. 2018). Thus, H_2_S may be mediating its toxicity preferentially through Ca_V_ 1.2 channels based on their relative ability to antagonize H_2_S-induced effects on SCO. These results also showed that H_2_S-induced neurotoxicity is not mediated through T-type receptors as blockage of T-type VGCC channels using ethosuximide failed to reverse H_2_S -suppressed SCO in PCN. Even though nifedipine showed efficacy in reversing H_2_S-induced suppression of SCO in vitro, this drug was reported to induce hypotension in vivo. H_2_S poisoning was also reported to induce hypotension (Baldelli et al. 1993) limiting the possible use of nifedipine as a drug for treating victims of H_2_S poisoning.

<u>T</u>ransient <u>R</u>eceptor <u>P</u>otential (TRP) channels mediate Store-operated Ca^2+^ (SOC) entry (Lopez et al. 2020) and have previously been implicated in H_2_S-induced pathophysiology (Ng et al. 2019; Pozsgai et al. 2019). Pre-exposure to 2-APB showed partial efficacy by preventing H_2_S-induced suppression of SOC in Phase 2. 2-APB, like nifedipine, failed to antagonize effects of H_2_S in Phase 1. Previously SCO was shown to require cyclic Ca^2+^ entry through the NMDA receptor in cerebellar neurons via activated P-type Ca^2+^ channel (Nunez et al. 1996). We have shown that H_2_S suppresses neuronal electrical activity in PCN, and that 10 μM 2-APB antagonized this, reversing it to a degree. TRP channels are a large group of channels with multiple sub-families, and diverse groups of proteins are involved (Samanta et al. 2018). 2-APB is a non-selective blocker of TRP channels and SOCE. 2-APB also antagonizes IP_3_R and store-operated Ca^2+^ channel in ER (Splettstoesser et al. 2007). Activation of SOC entry is triggered by the depletion of Ca^2+^ stores in ER (Bollimuntha et al. 2017). Dantrolene antagonizes ryanodine receptor in ER and inhibits calcium release. 2-APB may also play a role in inhibiting voltage-gated and calcium dependent potassium conductance and affecting on depolarization of membrane potential (Hagenston et al. 2009). Though promising, as a non-selective TRP blocker, 2-APB might induce off-target effects. More research is warranted to investigate the efficacy of 2-APB and to study the efficacy of other more selective TRP antagonists in acute H_2_S induced neurotoxicity.

Hydrogen sulfide is known to inhibit cytochrome c oxidase activity in mitochondria (Anantharam et al. 2017a) depleting ATP. It is plausible that H_2_S-induced depletion of ATP might lead to Ca^2+^ dysregulation via glutamate receptor. The NMDA receptor has been implicated in H_2_S-induced neurotoxicity because MK-801, an NMDA receptor inhibitor protects cerebellar granule neurons (Cheung et al. 2007; Garcia-Bereguiain et al. 2008). However, other investigators reported that H_2_S-induced neurotoxicity was not improved by use of NMDA receptor inhibitors (Kurokawa et al. 2011). In this study, MK-801 exacerbated H_2_S-induced suppression of SCO in PCN. This implies that H_2_S acting through NMDA receptors plays a role in regulating SCO in PCN through Ca^2+^ signaling. Pretreatment with perampanel also exacerbated H_2_S-induced suppression of SCO. Dravid and Murray showed that glutamate and AMPA receptors play a role in spontaneous SCO in neocortical neurons (Dravid and Murray 2004). Findings from this study are consistent with those observations and is further evidence that glutamate and AMPA receptors are involved in regulating SCO during H_2_S intoxication.

Hydrogen sulfide exposure induces many neurological dysfunctions including motor, behavioral, memory, and visual impairment, and induces neurological sequalae (Nam et al. 2004; Tvedt et al. 1991a) including neuronal cell death in select brain regions (Anantharam et al. 2018; Anantharam et al. 2017a; Anantharam et al. 2017b; Kim et al. 2018; Kim et al. 2019). Inhibition of the breathing center in the brainstem and/or prolonged coma have been linked to death and neurological complications (Guidotti 1994; Sonobe and Haouzi 2015). Calcium signaling is involved in mediating these effects. Preventing or reversing dysregulation of Ca^2+^ signaling may be one of the ways to treat acute H_2_S-induced neurotoxicity, and reduce mortality and morbidity.

## Conclusion

Hydrogen sulfide is a potent toxicant targeting the brain, lung, and heart. In this study, we examined brain, lung, and heart function in real-time during H_2_S exposure. H_2_S suppressed electrocerebral activity and disrupted breathing. Cardiac activity was comparatively less affected. This in vivo study provided direct evidence that it is the H_2_S-induced dysregulation of brain activity that triggers breathing challenges and not vice versa. It also showed that heart function is the last to fail of the 3 major target organs of H_2_S poisoning. We were also able to recapitulate the in vivo brain electrical activity using a simple in vitro model of PCN in two neuronal function assays FLIPR and MEA. Using the in vitro functional PCN model we showed that following a single acute H_2_S exposure, 3 phases of H_2_S-induced neurotoxicity were discernible using FLIPR and MEA. Phase 1 is characterized by hyperactivity. Phase 2 is characterized by suppressed electrical activity. Phase 3 is characterized by recovery. H_2_S consistently induced phase-specific changes in peak count, peak width, peak rise time, and peak decay time. Using these parameters, we showed that nifedipine, an L-type VGCC inhibitor antagonized H_2_S-induced suppression of SCO in PCN. Surprisingly, nimodipine, another L-type VGCC antagonist, failed. Blockage of SOC entry by 2-APB also prevented H_2_S-induced suppression of SCO and neuronal electrical activity and was protective of the PCN. NMDA and AMPA receptors were shown to be involved in H_2_S-induced regulation of SCO. For example, pre-treatment with perampanel effectively antagonized effects of H_2_S in Phase 1. However, like MK- 801, it worsened the effects of H_2_S in Phase 2, further suppressing SCO. Notably, we have developed a rapid in vitro system consisting of FLIPR and MEA to study the role of various Ca^2+^ channels in H_2_S-induced neurotoxicity. Used in conjunction with ImagExpress^®^ high content imaging, we can also assess the neuroprotective efficacy of test articles and drug candidates to treat acute H_2_S-induced neurotoxicity. Drugs that can normalize H_2_S-induced Ca^2+^ dysregulation may have therapeutic potential in reducing H_2_S-induced mortality and morbidity. We plan to use this in vitro system to further understand the role of calcium in H_2_S induced neurotoxicity and for potential discovery of novel and repurposed drugs for treatment of acute H_2_S poisoning to reduce mortality and/or morbidity.

---

**Supple Fig. 1.**
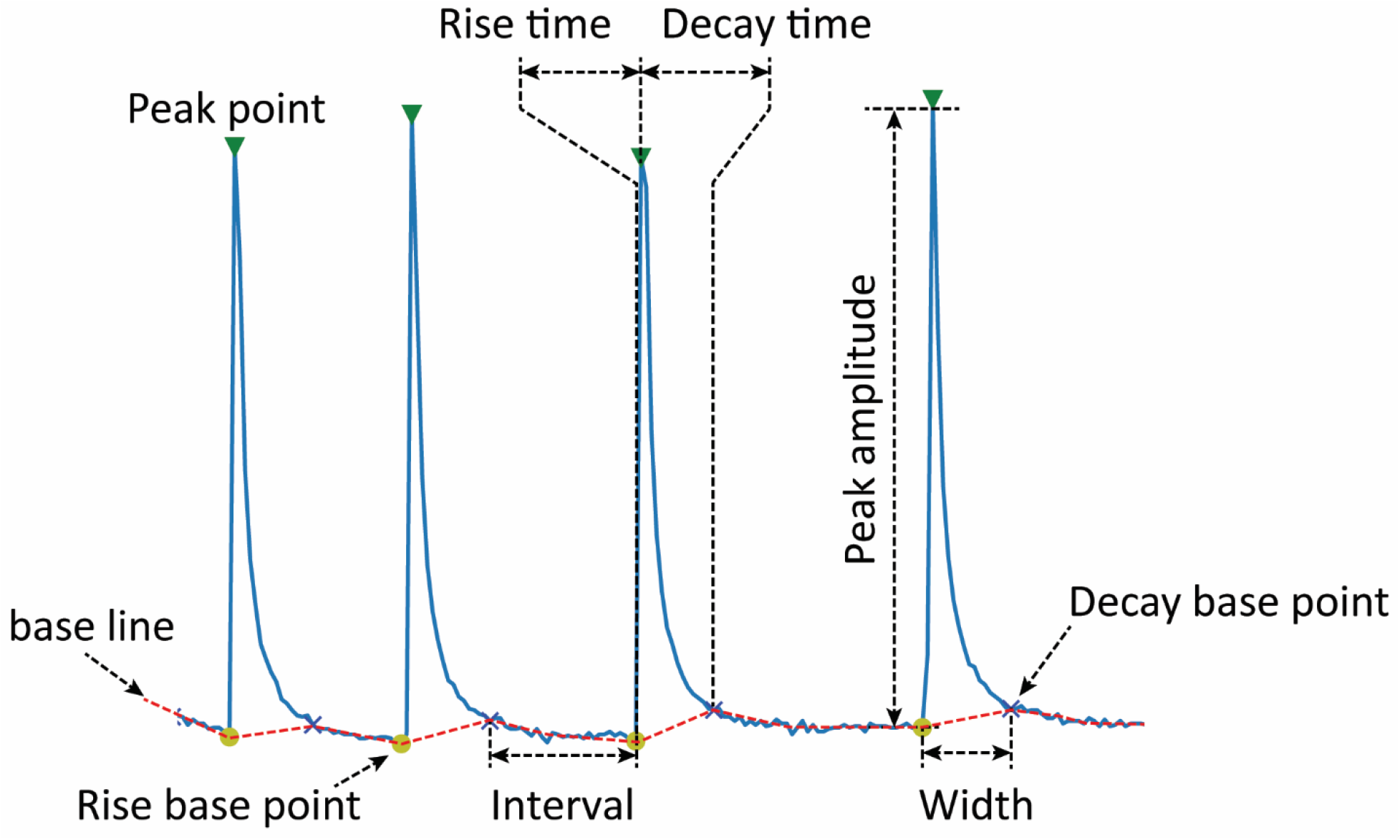
Typical FLIPR output of synchronous calcium oscillation (SCO) in normal primary cortical neurons. Measurements of variations in this typical FLIPR output was used to assess the effects of H_2_S on Ca^2+^ channels and the impact of pharmacological probes on SCO.

**Supplementary Fig. 2.**
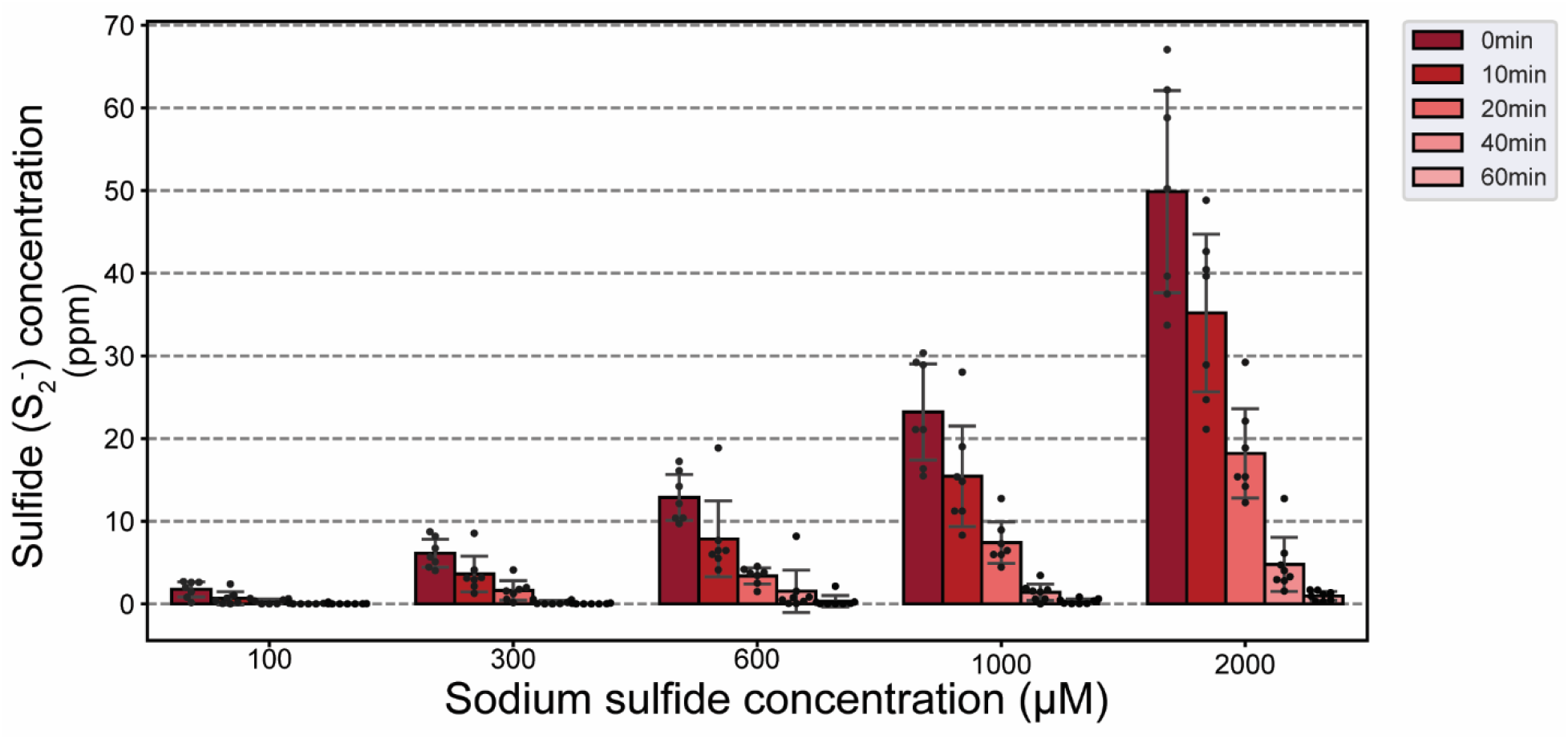
Summary of dose and time-dependent changes in H_2_S concentrations following addition of Na_2_S, a chemical donor of H_2_S. A series of concentrations of Na_2_S were dissolved in SCO imaging solution, and sulfide concentrations measured using a sulfide microelectrode in the solution over a 60 min time period. From this experiment, we chose to use 2000 μM Na_2_S which generated 34 ppm H_2_S measured as sulfide for all our definitive in vitro experiments.

